# Context-dependence in the symbiosis between *Dictyostelium discoideum* and *Paraburkholderia*

**DOI:** 10.1101/2021.09.17.459779

**Authors:** Trey J. Scott, David C. Queller, Joan E. Strassmann

**Affiliations:** Department of Biology, Washington University in St. Louis

**Keywords:** symbiosis, context-dependence, competition, bet-hedging, *Dictyostelium discoideum*, *Paraburkholderia*

## Abstract

Symbiotic interactions change with environmental context. Measuring these context-dependent effects in hosts and symbionts is critical to determining the nature of symbiotic interactions. We investigated context-dependence in the symbiosis between social amoeba hosts and their inedible *Paraburkholderia* bacterial symbionts, where the context is the abundance of host food bacteria. *Paraburkholderia* have been shown to harm hosts dispersed to food-rich environments, but aid hosts dispersed to food-poor environments by allowing hosts to carry food bacteria. Through measuring symbiont density and host spore production, we show that this food context matters in three other ways. First, it matters for symbionts, who suffer a greater cost from competition with food bacteria in the food-rich context. Second, it matters for host-symbiont conflict, changing how symbiont density negatively impacts host spore production. Third, data-based simulations show that symbiosis often provides a long-term fitness advantage for hosts after rounds of growth and dispersal in variable food-contexts, especially when conditions are harsh with little food. These results show how food context can have many consequences for the *Dictyostelium-Paraburkholderia* symbiosis and that both sides can frequently benefit.

**Impact Statement:** Many organisms form symbiotic relationships with other species. These symbioses often exhibit context-dependence, where the sign or magnitude of one partner’s effect on the other will change in different environments. Context-dependent effects make it difficult to assign interactions to categories like mutualisms or antagonisms because they involve both benefits and costs depending on the environment. However, in some cases, accounting for context-dependence can clarify an interaction so that it more easily fits a mutualism or antagonism. We investigated context-dependence using the symbiosis between *Dictyostelium discoideum* and two symbiotic *Paraburkholderia* species. In this symbiosis, *Paraburkholderia* bacteria allow hosts to carry food bacteria to food-poor contexts, where hosts rarely survive without food, but reduce host fitness in the more hospitable food-rich contexts. The effect of food context on P*araburkholderia* symbionts is unknown. We show that *Paraburkholderia* symbionts are also affected by this context, through facing reduced competition after being dispersed by hosts to food-poor contexts. We also identify a new way that symbionts affect hosts, where symbiont density reduces host fitness, but less so in food-poor contexts. Finally, we use simulations to show that infected hosts benefit in the long-term across variable food contexts, especially in the harshest environments with little food. These results show that context-dependence in symbiosis can have many consequences for hosts and symbionts, though in general for *D. discoideum* and *Paraburkholderia*, both are likely to benefit.

## Introduction

Context-dependence, where the environment can change the sign or magnitude of one partner’s effect on the other, is common in symbioses (Bronstein, 1994; Thompson, 1994; Chamberlain *et al*., 2014). These context-dependent effects on partners can be crucial to understanding the nature of symbiotic interactions (Keeling & McCutcheon, 2017; Iwai, 2019). For example, in the symbioses between *Paramecium bursaria* hosts and their *Chlorella* endosymbionts, hosts benefitted from symbiosis in light environments, but were harmed in the dark. For *Chlorella*, the effects of symbiosis were negative in co-culture, indicating that hosts exploited their endosymbionts for the benefits hosts receive in light conditions (Lowe *et al*., 2016). However, in the context of an environment with a *Chlorella* competitor, hosts benefited their symbionts by eating these competitors (Iwai, 2019). This example illustrates that understanding how partners affect each other across multiple contexts can change our view of the interaction, sometimes from one of exploitation to one of mutual benefit.

Context-dependence is important in the lifecycle of the social amoeba *Dictyostelium discoideum*. Amoebae need edible bacteria to grow and proliferate (Raper, 1937), but the abundance (Young, 2004; Vos *et al*., 2013) and quality (Kuserk, 1980; Brock *et al*., 2018) of food bacteria in the soil is known to vary. This results in a patchy environment where some patches are food-rich and other patches are food-poor. In response to starvation, amoebae aggregate and form a multicellular fruiting body to disperse resistant spores to new environments (smith *et al*., 2014). The patchy soil environment is considered an important selection pressure for this fruiting body structure (Bonner, 1982; Kessin, 2001).

*D. discoideum* interacts with three species of mostly inedible *Paraburkholderia* bacterial symbionts — *P. agricolaris, P. hayleyella*, and *P. bonniae* (Brock *et al*., 2020). Throughout this paper, we will use “*Paraburkholderia*” or “symbionts” as shorthand for the three symbiotic *Paraburkholderia* species. Hosts infected with *Paraburkholderia* have been isolated from multiple locations in the United States, with around 25% of screened hosts being infected by at least one species (Haselkorn *et al*., 2019). *Paraburkholderia* are able to enter and live inside *D. discoideum* cells and spores, but can also proliferate, albeit sometimes only slowly, without their hosts (DiSalvo *et al*., 2015; Shu *et al*., 2018a; Brock *et al*., 2020) unlike the obligate endosymbionts that are also found in *D. discoideum* (Haselkorn *et al*., 2021). There is some evidence consistent with coevolution between hosts and symbionts (Brock *et al*., 2016; Shu *et al*., 2018a; Garcia *et al*., 2019; Brock *et al*., 2020). For example, host clones naturally infected with *P. hayleyella* are harmed less by infection with this symbiont than host clones that were not infected in the wild (Shu *et al*., 2018a) indicating that *P. hayleyella* hosts have adaptations favoring symbiosis. Symbionts also have the ability to move towards hosts (Shu *et al*., 2018b), suggesting that being able to find hosts is beneficial.

The symbiosis with *Paraburkholderia* bacteria impacts the growth and proliferation of *D. discoideum*. Having symbionts allows hosts to carry food bacteria (and inedible *Paraburkholderia*) inside the spore-containing part of fruiting bodies called the sorus (DiSalvo *et al*., 2015). Whether this novel trait is advantageous or not depends on the presence of food bacteria after dispersal. When food is abundant, having symbionts can be costly, as shown by infected amoebae producing fewer spores than uninfected amoebae (Brock *et al*., 2011; DiSalvo *et al*., 2015). In food-poor environments, the cost of having *Paraburkholderia* is compensated by hosts gaining the ability to carry food bacteria in dispersing spores. This allows amoebae to disperse and grow where they ordinarily could not (Brock *et al*., 2011; DiSalvo *et al*., 2015). These context-dependent effects on the host could be extremely important in the natural soil environment, where food-poor patches arise frequently (Kessin, 2001).

Less is known about how symbionts are affected across food contexts. Gaining the ability to disperse to new locations may be a major reason for symbionts to seek out social amoeba hosts (Garcia & Gerardo, 2014), but could also make the context of host food bacteria in the new environment important for symbionts. A benefit from being dispersed to patches with few bacteria could be that symbionts face reduced competition. If few bacteria are present, symbionts will mostly compete with food bacteria that were also carried in the sorus. This should be a relatively low competition situation because symbionts outnumber food bacteria in sori (Khojandi *et al*., 2019). Having few competitors should advantage symbionts while environments with plentiful bacteria could strongly limit symbiont growth because of their relatively slow growth rates, at least as measured in the lab (Brock *et al*., 2020). We will use “food-rich” and “food-poor” to describe newly colonized patches with many and few bacteria, respectively, of the sort edible by *D. discoideum*. These categories reflect the relationship to *D. discoideum* and could be called high and low competition in terms of their effect for *Paraburkholderia*.

It is unclear how the number of extracellular *Paraburkholderia* in the environment impacts hosts since previous studies have focused on intracellular *Paraburkholderia* (Shu *et al*., 2018b; Miller *et al*., 2020). When they are outside the amoebae, *Paraburkholderia* could affect *D. discoideum* fitness through interactions with food bacteria perhaps by reducing the amount of food for hosts through competition or by releasing diffusible toxins that affect amoebae. Thus host food context could also affect the relationship between symbiont density and host spore production.

The fitness effects of symbiosis for hosts have been tested only in food-poor and food-rich contexts individually. The benefits of symbiosis could pay out over the long-term across different food contexts in the soil. Growth rates in temporally variable contexts are best captured by geometric mean fitness rather than arithmetic mean fitness because only the geometric mean captures the lasting effects of periods of low fitness (Sæther & Engen, 2015). Ignoring geometric means can lead to incorrect assessments of the adaptive value of strategies in variable environments. One example of an adaptation that is only apparent from geometric mean fitness measures are bet-hedging phenotypes, where organisms adapt to uncertain environments by avoiding the worst effects of harsh contexts while being suboptimal in more favorable contexts (Slatkin, 1974; Philippi & Seger, 1989; Starrfelt & Kokko, 2012). This lowers the variance in fitness across time and results in higher geometric mean fitness at the expense of lower arithmetic mean fitness.

Bet-hedging is suspected to play a role in explaining observations of disadvantageous partnerships in plant-fungus mutualisms (Lekberg & Koide, 2014; Veresoglou *et al*., 2021). If bet-hedging occurs, short-term costs are acceptable if partnerships increase geometric mean fitness. Alternatively, symbiosis could increase both geometric and arithmetic mean fitness across contexts without the need for bet-hedging. In this case, the benefits of symbiosis simply outweigh the costs as in more traditional descriptions of symbioses (Douglas, 2010). However, these alternatives have not been tested in detail.

To understand context-dependence in the symbiosis between amoebae and *Paraburkholderia*, we used *D. discoideum* infected with either *P. agricolaris* or *P. hayleyella —* the two most common and best-studied species of *D. discoideum* symbionts. We investigate whether *Paraburkholderia* benefit from reduced interspecific competition when dispersed to food-poor contexts, how symbiont density and food context impact host spore production, and whether symbiosis is beneficial for hosts when food conditions vary.

## Methods

To understand the effects of symbiosis across food contexts, we used 4 naturally uninfected *D. discoideum* clones, 4 clones naturally infected with *P. agricolaris*, and 4 clones naturally infected with *P. hayleyella* (Figure 1). We cured the infected clones and re-infected them with their native symbionts to standardize infection density. Uninfected clones were left uninfected, but were otherwise treated the same as infected clones. This resulted in three host infection conditions: uninfected, infected with *P. agricolaris*, and infected with *P. hayleyella*. To mimic natural dispersal, we collected sori and transferred them to food-rich (with additional *K. pneumoniae* bacteria) or food-poor (KK2 buffer with no *K. pneumoniae*) nutrient plates. Bacteria appear on food-poor plates only if transferred sori contain bacteria, as expected for infected samples. We grew replicate experimental sets involving all conditions beginning on two separate dates, July 13 and 22, 2020, and followed up with additional experiments (see results) beginning on January 26 and April 23, 2021.

**Figure 1:**
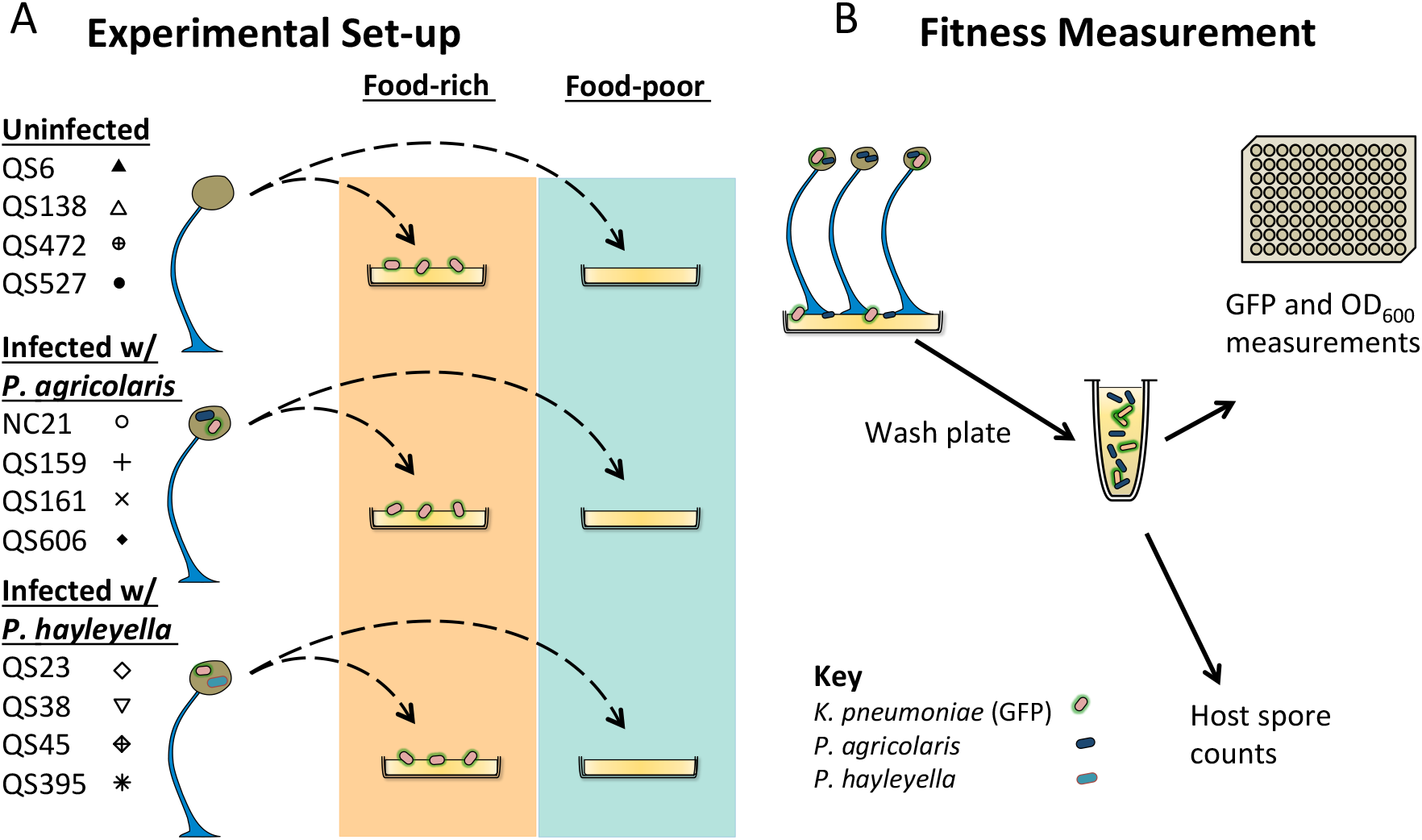
Schematic of experimental design. (A) Uninfected and infected *D. discoideum* fruiting bodies are collected and plated on food-rich and food-poor plates (after one passage on GFP-expressing *K. pneumoniae* food bacteria). These plates are grown for six days and then washed for bacterial measurement and spore counting (B). Bacteria are measured by calculating GFP fluorescence and optical density (see Methods). Host spore production is measured from washed plates.

### Paraburkholderia isolation

To isolate *Paraburkholderia* from their hosts, we grew wild collected *D. discoideum* clones on SM/5 plates (2 g glucose (Fisher Scientific), 2 g Bacto Peptone (Oxoid), 2 g yeast extract (Oxoid), 0.2 g MgSO_4_ * 7H_2_O (Fisher Scientific), 1.9 g KH_2_PO_4_ (Sigma-Aldrich), 1 g K_2_HPO_4_ (Fisher Scientific), and 15 g agar (Fisher Scientific) per liter). Wild *D. discoideum* clones were grown with *K. pneumoniae* food bacteria that were suspended in KK2 buffer (2.25 g KH_2_PO_4_ (Sigma-Aldrich) and 0.67 g K_2_HPO_4_ (Fisher Scientific) per liter). After wild clones completed the social cycle (feeding, starvation, and fruiting body formation), we collected sori with pipette tips and placed them on SM/5 plates. We allowed the bacteria and amoebae contained within to proliferate and then streaked out the resulting bacteria to get single colonies.

### Paraburkholderia removal

To generate uninfected clones, we treated infected *D. discoideum* clones with antibiotics by plating on 30 μg/mL tetracycline SM/5 plates with 200 μL of 1.5 optical density (OD_600_) tetracycline-resistant *K. pneumoniae* suspended in KK2 buffer. After passage on SM/5 plates without tetracycline to let the amoebae recover from any effects of the antibiotic, we collected single sori with a pipette tip and placed ten of them in different locations on SM/5 plates to confirm that we had successfully removed the bacteria. If bacteria are present, these spot tests will show bacterial growth and *Dictyostelium* proliferation as the spores hatch and eat the bacteria (Brock *et al*., 2011). Without bacteria, amoebae cannot proliferate and the spot will stay blank. We considered a clone to be cured if no bacteria showed up in spot tests. We similarly treated naturally uninfected hosts with tetracycline to control for any effect of curing on our results.

### Paraburkholderia re-infection

We re-infected cured *D. discoideum* clones with their native *Paraburkholderia* isolates by plating 200 μL 2×10^5^ spores with 200 μL of 0.1% *Paraburkholderia* solution. This solution consisted of 1.5 OD_600_ *Paraburkholderia* and 1.5 OD_600_ *K. pneumoniae* in a 1:1000 ratio. To confirm re-infection (and also successful isolation), we performed spot tests as above, where successful re-infection was inferred when bacteria grew on 8 or more spots out of the 10 we put down.

### Artificial dispersal to food-rich and food-poor plates

To obtain sori to transfer to food-rich and food-poor plates, we started by growing *D. discoideum* clones from frozen stock, as described above, on 200 μL 1.5 OD *K. pneumoniae* expressing green fluorescent protein (GFP). We obtained GFP-expressing *K. pneumoniae* (strain ID DBS0349837) from the Dicty Stock Center at dictyBase (Fey *et al*., 2013). This initial growth period is to remove freezer effects and ensure that food bacteria that are carried to new plates are GFP-expressing since stocks were fed non-GFP bacteria before freezing. After six days of growth, we used pipette tips to collect sori from mature fruiting bodies. We counted spores using a hemocytometer and diluted spores to a concentration of 2×10^5^ per mL, and then plated them on plates with (food-rich) or without (food-poor) an additional 200 μL of the GFP-expressing food bacterium *K. pneumoniae* (Figure 1). In order to survive on food-poor plates, the host must carry food bacteria from the previous plate. We grew food-rich and food-poor plates for six days unless otherwise stated, enough time for mature fruiting bodies to form.

### Measurement of bacteria density

To measure *Paraburkholderia* density, we measured the quantity of bacteria left on plates after *D. discoideum* formed fruiting bodies. We first collected plate contents by washing plates with 15 mL of KK2 buffer. To remove fruiting bodies and bacteria associated with fruiting bodies, we centrifuged wash solutions for three minutes at 13000 rpm. We measured bacteria using optical density measured at 600 nm (OD_600_), a frequency at which bacteria commonly scatter light. Because the OD_600_ is due to both *Paraburkholderia* and *K. pneumoniae*, we used GFP fluorescence measurements (with an excitation wavelength of 485 and emission wavelength of 515 nm) and a standard curve relating *K. pneumoniae* fluorescence to its OD_600_ to subtract out the component due to GFP-expressing *K. pneumoniae*. Both OD_600_ and fluorescence measures were performed in a 96 well plate with a Tecan Infinite 200 Pro microplate reader.

To validate our standard curve, we compared predicted OD_600_ of *P. agricolaris* and *K. pneumoniae* to colony forming unit (CFU) counts from the same samples. Linear regression revealed that predicted OD_600_ measurements explained most of the variation in CFUs, showing that our assay is reliable (Figure S1). We also checked our standard curve for significant quadratic terms, which can cause measurement errors when combining OD_600_ and fluorescence measures at high densities (Meyers *et al*., 2018), but our curve did not have a significant quadratic term.

### Host spore production

Spore production is a standard fitness measure in *D. discoideum* (Buttery *et al*., 2009; Hall *et al*., 2013; Gruenheit *et al*., 2017). To measure host spore production, we estimated spore concentration in the supernatants from washed plates using a hemocytometer. We then calculated the total number of spores per plate by multiplying by the volume of wash solution.

### Spore production simulations

To test whether infected hosts benefit across variable food contexts, we simulated rounds of growth and dispersal across soil patches with different probabilities of having food bacteria. We separately modeled three host phenotypes: (1) uninfected, (2) infected with *P. agricolaris*, and (3) infected with *P. hayleyella*. Co-infections are possible, but are rare in nature (Haselkorn *et al*., 2019) so we exclude them from our analysis.

We assumed that environments consisted of 100 discrete soil patches. Patches were either food-poor or food-rich (we investigated continuous amounts of food and found similar results; see Supplemental file 1). Food-poor patches at time *t* were drawn from a binomial distribution with probability *p_t_*. Food-rich patches were drawn with probabity 1-*p_t_*. To allow temporal variation, the value of *p_t_* in each generation was drawn from a beta distribution with mean *p* and variance *v_temp_*. High values of v_temp_ resulted in more temporally variable environments. For low values, most of the variation was spatial.

Initially all patches were colonized. Each patch produced a number of spores, drawn from the distribution of our empirical spore production values, according to whether it was a food-rich or food-poor patch. To model costs, we penalized host spore production in food rich environments by reducing spore production by a percentage *c*. When *c* is 0, we modeled the scenario observed in this study, with no infection cost. We did not detect a cost of infection in food-rich contexts, but numerous other studies have documented this cost (Brock *et al*., 2011; DiSalvo *et al*., 2015; Miller *et al*., 2020). It is likely that we did not detect a cost because we infected hosts with fewer *Paraburkholderia*. Because these costs have been demonstrated repeatedly in other studies and because of the importance of costs to bet-hedging (Lekberg & Koide, 2014; Veresoglou *et al*., 2021), we included them as a variable. We summed cost-adjusted spore production values to get the total spore production across all patches. This is divided by 2×10^5^, a rough estimate of the number of spores in a typical sorus, to get the total number of sori, which we are assuming to be the dispersal unit.

The global pool of sori is used to seed the next round. New patches are assumed to be empty and dispersal is assumed to be global such that sori from one patch can disperse to any other patch with equal probability. Dispersal is likely efficient in *D. discoideum* as sori can be dispersed long distances by arthropods (smith *et al*., 2014) and possibly even by birds (Suthers, 1985). Each sorus is randomly assigned to a patch and it successfully colonizes that patch *g*% of the time. Because the value of *g* for natural hosts is unknown, we investigated three values of *g* (50%, 5%, and %0.5) that range from cases where there are many more sori successfully establishing than available patches to cases where each patch produces around one sorus. We assumed that patches colonized by multiple sori were the same as singly colonized patches for the purposes of determining their subsequent spore production. Some patches may remain unfilled (though this is unlikely when g = 50%). We also assume that infection status is not associated with different rates of colonization as those differences are better captured by our empirical spore production values, which will include differences in growth efficiency or spore germination rate.

We vary the average probability of food-poor patches *p* from 0.1 to 0.9 and simulate four different cost regimes reflecting variation found in different *Paraburkholderia* isolates (Miller *et al*., 2020). We simulated dispersal to new patches for 100 rounds of growth and dispersal using 100 replicates for each combination of *p*, *v_temp_*, and *c* for each phenotype. Within each replicate, all three phenotypes experience the same environment. At the end of the 100 rounds, we calculated the total spore production per round and calculated geometric and arithmetic mean spore production from these values across the 100 rounds. Within each replicate, we determined whether infected hosts had higher geometric or arithmetic mean fitness for each individual simulation and whether any phenotype went extinct.

We assigned outcomes for each parameter combination by calculating the frequency that infected or uninfected hosts had higher geometric mean fitness or arithmetic mean fitness. Infected and uninfected hosts were assigned as winners if they had higher geometric mean fitness in 75% of replicates. We assigned an outcome as bet-hedging when infected hosts won and more than half of the winning replicates did so with lower arithmetic mean fitness. Extinctions occurred in some simulations and were treated as a distinct outcome. We assigned mixed outcomes when neither infected nor uninfected hosts were able to have higher geometric mean fitness in 75% of replicates. Some mixed outcomes involved individual replicates where infected hosts were found to bet-hedge.

### Statistical Methods

We performed statistics in R version 3.6.3 (R Core Team, 2020). To compare bacteria density and spore production, we used linear mixed models (LMM) with the *lme* function in the nlme package (Pinheiro & Bates, 2006). To account for random variation from replicate clones and effects of dates when experiments were performed, we included clone and the date the experiment was performed — along with each variable on its own — as random effects. To select the best model of random effects, we used AICc, a sample-size-corrected measure of model fit that balances predictive ability and model complexity (Burnham & Anderson, 2004). Many of our models showed different variances between treatments. To account for these differences in variance, we weighted models with the *varIdent* function in nlme (Pinheiro & Bates, 2006). We used the emmeans package (Lenth *et al*., 2018) to perform contrasts.

To understand how *Paraburkholderia* density affects host spore production across food conditions, we fit a LMM using only infected hosts that included symbiont density leftover on plates and whether the plate was food-rich or food-poor, along with the interaction between these variables. We included random effects for clone, date, and both crossed effects and selected the best random effect structure with AICc. We determined whether the interaction was important by comparing AICc of the model including the interaction with models including the other variables but lacking the interaction.

## Results

### Paraburkholderia dispersed by Dictyostelium sori have lower growth when host food bacteria are abundant

The context of a food-poor environment is known to be important for *D. discoideum* hosts. It is not known how *Paraburkholderia* are affected by this same context, but reduced competition with food bacteria seems likely. We tested this by growing infected sorus contents on food-poor and food-rich nutrient plates and measuring the density of *Paraburkholderia* after *D. discoideum* fruiting body formation (Figure 1). After infected hosts formed fruiting bodies, *Paraburkholderia* densities were lowest in food-rich conditions (Figure 2A), as expected if they compete with food bacteria. There was around five times more *P. agricolaris* on food-poor than food-rich plates (LMM, p < 0.001). *P. hayleyella* growth was higher in food-poor conditions than food-rich, but this difference was not significant after 6 days (LMM, p = 0.416). Because *P. hayleyella* grows slowly, we performed two more experiments with *P. hayleyella* with 8 and 12-day growth periods (Figure 2B). Allowing for longer incubations did not result in significantly higher density of *P. hayleyella* (LMM, p = 0.633), suggesting that *P. hayleyella* reach their maximum density at or before 6 days, but including these additional experiments gave us enough power to find a significant increase in *P. hayleyella* density in food-poor conditions relative to food-rich (LMM, p = 0.027). These results show that symbiont density is context-dependent.

**Figure 2:**
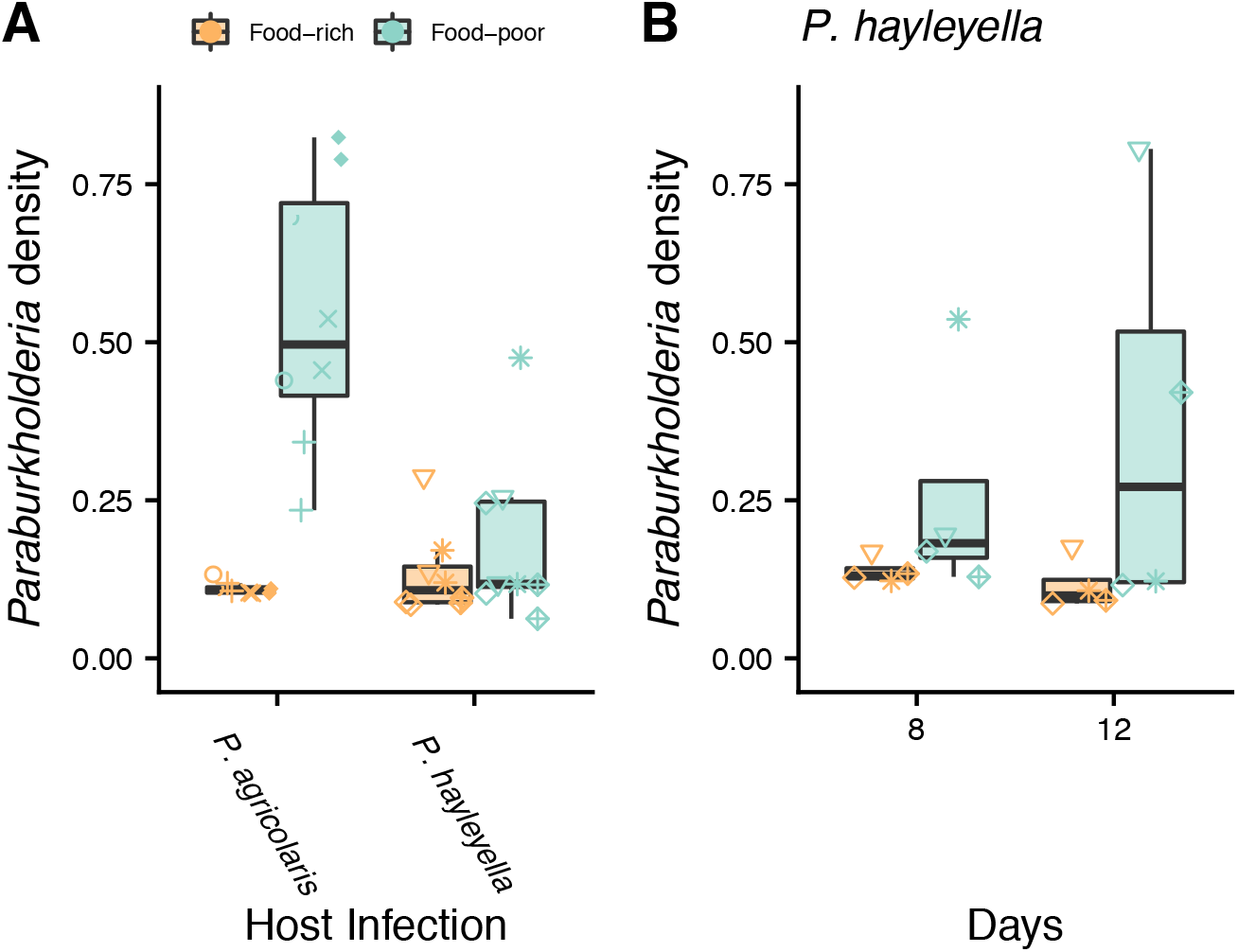
More *Paraburkholderia* were recovered from plates after fruiting body formation from food-poor plates (those that had not received additional *K. pneumoniae*). (A) *Paraburkholderia* density after 6 days. (B) *P. hayleyella* density after 8 and 12 days. Point shapes show individual clones (see Figure 1).

### Higher symbiont density harms hosts, but less so in food-poor contexts

The host-food context may affect the relationship between symbiont density and host spore production and therefore the degree of conflict or cooperation between them. To investigate this, we also measured total host spore production from plates where we measured the growth of *Paraburkholderia* symbionts (Figure 1). We used uninfected hosts as a baseline for fitness without symbionts. We confirmed prior studies (Brock *et al*., 2011; DiSalvo *et al*., 2015) showing that infected hosts could carry food bacteria and proliferate on food-poor plates, while uninfected host could not (Figure 3A). Surprisingly, we did not observe a cost of being infected in food-rich conditions (p > 0.5 for both species) which has been seen in previous studies (Brock *et al*., 2011; DiSalvo *et al*., 2015; Shu *et al*., 2018a). This is likely a result of our lower infection dosage of 0.1%.

**Figure 3:**
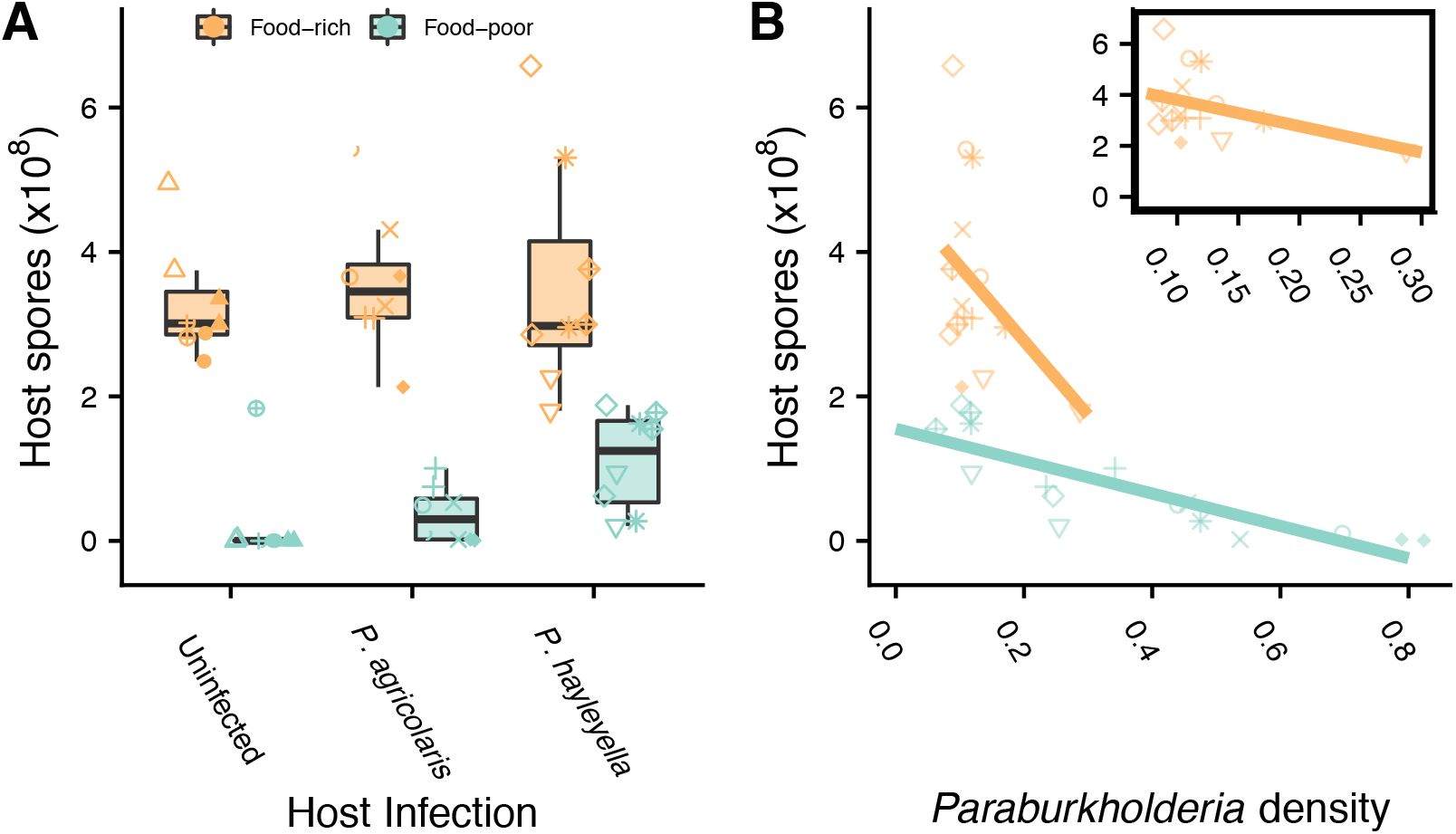
Effects of *Paraburkholderia* infection and density on host spore production. (A) Spore production of hosts from food-rich and food-poor plates for uninfected, *P. agricolaris* infected, and *P. hayleyella* infected hosts. (B) Interaction between measured *Paraburkholderia* density (OD_600_) and food environment on host spore production. This interaction model explained 95% of the variance in spore production. Inset shows food-rich results on smaller scale. Point shapes show individual clones (see Figure 1).

While having some symbionts is essential for hosts to be able to carry food and survive in food-poor conditions, higher symbiont densities may nevertheless harm hosts, perhaps in ways that depend on food context. We found that larger populations of symbionts as measured by OD_600_ were associated with lower host spore production, but this harm was reduced in food-poor conditions. Lower host spore production was associated with being in a food-poor environment (β_food-poor_ = −3.283, se = 0.853) and symbiont density (β_density_ = −10.317, se = 6.364), but the interaction between food scarcity and symbiont density showed that the harmful effect of higher symbiont densities was lessened on food-poor plates (β_food-poor*density_ = 8.078, se = 6.381; Figure 3B). These results indicate that symbiont density may come at the expense of host spore production, but that this cost decreases in food-poor environments.

### Symbiosis is often beneficial for hosts across variable contexts

Because symbiosis helps hosts in food-poor contexts, we hypothesized that infected hosts would gain a long-term benefit across contexts compared to uninfected hosts. If infected hosts increased their geometric mean fitness at the expense of arithmetic mean fitness, infected hosts could even gain a bet-hedging advantage. We modeled this by using our empirical spore production values to simulate 100 rounds of growth and dispersal across environments where the number of food-poor patches was determined by the mean frequency (*p*) and the temporal variance (*v_temp_*; more detail can be found in the methods). Because the natural conditions of this symbiosis are mostly unknown, we simulate a wide-range of parameter space to determine which conditions favor symbiosis. The supplement includes animations of representative simulations.

We first describe the results when dispersing sori successfully colonize new patches 5% of the time. When there was no cost of infection, we found that infected hosts were favored in every condition we tested (Figure 4; blue). We also simulated costs of infection because those have been found in other studies (Brock *et al*., 2011; DiSalvo *et al*., 2015; Miller *et al*., 2020). As the cost of infection in food-rich contexts increased, infected hosts were favored in the most food-poor environments while uninfected hosts were favored when food was abundant (Figure 4; orange). *P. hayleyella* was favored across more environments than *P. agricolaris*.

**Figure 4:**
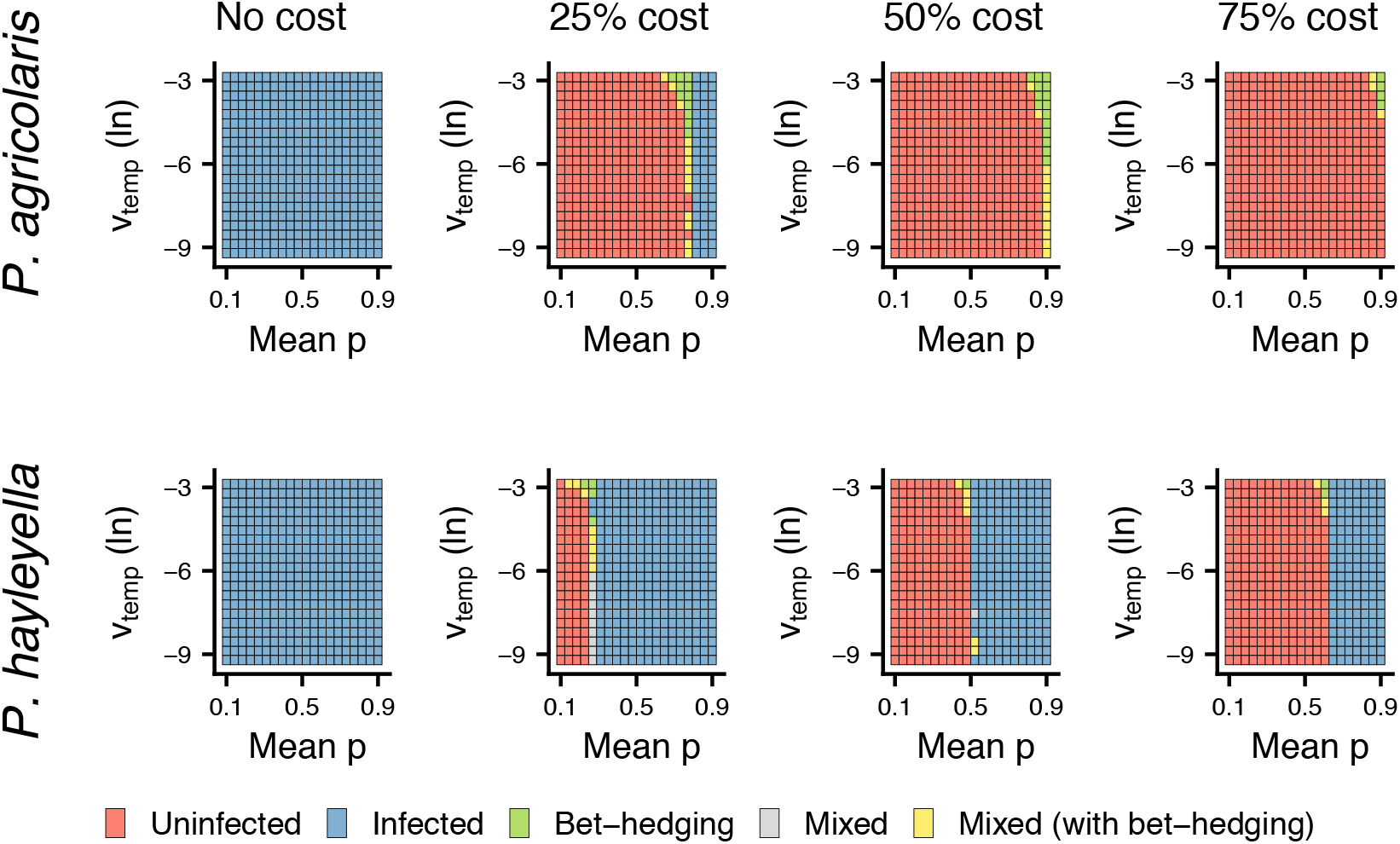
Benefits of symbiosis depends on variation in food availability and fitness costs. Winning phenotypes of *P. agricolaris* (top) and *P. hayleyella* (bottom) relative to uninfected for different costs of infection with a 5% probability of colonization. Orange shows when uninfected hosts have higher arithmetic and geometric mean spore production; blue shows when infected hosts have higher arithmetic and geometric mean spore production; green shows when arithmetic fitness is reduced for higher geometric mean fitness (bet-hedging); gray shows areas where both infection strategies can win; yellow shows where both strategies can win and where infected hosts bet-hedge.

Bet-hedging in this symbiosis appears to be rare (Figure 4; green and yellow). Infected hosts had a bet-hedging advantage when costs were added and food was intermediately rare. More temporally variable environments had a weak effect on increasing the likelihood of bet-hedging.

When dispersing sori successfully colonize new patches 50% of the time (each patch produces enough sori to completely fill the patches in the next generation), we found similar results (Figure S2). When only 0.5% were successful (each patch may only produce one or two sori for dispersal), we again found similar results except in the most food-poor conditions, where both uninfected and *P. agricolaris* infected hosts tended to go extinct (Figure S3A). *P. hayleyella* infected hosts were able to survive in these food-poor contexts (Figure S3B).

The natural environment of hosts is unlikely to involve food patches that are binary. Variation in the environment is also often auto-correlated, with the state of the environment at one time more often resembling the state of the environment in the near future (Ruokolainen *et al*., 2009). To determine whether our results were robust to variable environments with continuous food and temporal correlations, we ran additional simulations (described in detail in Supplemental file 1) where the amount of food varied from 0 to 1 (Figure S4) depending on a continuous resource that allowed us to tune autocorrelations (Figure S5). These additional simulations broadly supported our conclusions from the simpler simulations (Figure S6-8).

## Discussion

Our results show how the context of host food abundance affects the *Dictyostelium*-*Paraburkholderia* symbiosis beyond the previously demonstrated advantage to hosts when food is rare (Brock *et al*., 2011). First, we found evidence that both *Paraburkholderia* species benefit from reduced competition when they are carried to food-poor environments (Figure 2). Second, symbiont density negatively affected host spore production, but symbionts harmed hosts less in food-poor conditions (Figure 3B). Third, infected hosts had an advantage over uninfected hosts in simulations when food conditions were harsh or when the cost of symbiosis was low (Figure 4).

Our finding that symbionts had higher growth when dispersed to food-poor contexts shows that *Paraburkholderia* symbionts experience parallel context-dependence as hosts. These results highlight the importance of context-dependence for both partners. *Paraburkholderia* may benefit from reduced competition when hosts bring them to food-poor environments because symbionts interact with fewer competitors or because hosts eat competitors. This, together with our finding that hosts can benefit across contexts, points to a relationship of mutual benefit in this symbiosis. Our results also fit with other findings of competitive benefits for symbionts (Iwai, 2019). Other benefits of symbiosis for *Paraburkholderia* remain to be tested.

Competition between symbionts and food bacteria may also be responsible for the context-dependent effects of symbiont density on host spore production. Our spore production results showed that higher symbiont densities resulted in lower host spore production, indicating that symbionts are harmful to hosts. However, higher symbiont densities are less harmful in food-poor conditions when competition is lower (Figure 3B). The reduced harm for hosts could be the result of less antagonism between bacteria, which results in less collateral damage to amoebae through secreted toxins or other competitive interactions between food bacteria and symbionts. The generality of our results is limited somewhat by only using one species of food bacteria. While using a single food bacterium is more experimentally tractable, amoebae encounter multiple bacteria species in their natural environments (Brock *et al*., 2018). Different species, or combinations of species, could change competition with symbionts and affect host spore production in different ways.

Symbiosis benefits amoeba hosts by giving hosts the ability to carry food to food-poor contexts (Brock *et al*., 2011; DiSalvo *et al*., 2015). Using simulations, we showed that this ability resulted in higher fitness across variable contexts when costs were low and food was rare (Figure 4). Under conditions with plentiful food and high costs, being uninfected was advantageous. In nature, about 25% of clones are infected (Haselkorn *et al*., 2019), suggesting that symbiosis is not universally favored. This indicates that our finding of no cost to hosts in the symbiosis may be unrepresentative of many natural infections. On the other hand, a 25% infection rate is high if the symbiosis is generally harmful. This indicates that the prevalence of symbiosis could reflect a balance of forces where *D. discoideum* is not strongly selected to fight *Paraburkholderia* infection in a geographic mosaic of coevolution (Thompson, 1994). Unfortunately, the natural conditions of this symbiosis are the biggest unknowns in this system as it is difficult to study this symbiosis, and microbes more generally (Kraemer & Boynton, 2017), in nature.

Hosts could also benefit across contexts through bet-hedging, where geometric mean fitness trades off with arithmetic mean fitness (Seger & Brockmann, 1987). It is suspected that costly symbioses may be able to evolve because they are advantageous over the long-term even if they are not advantageous in the short term (Lekberg & Koide, 2014; Veresoglou *et al*., 2021). We found that bet-hedging was rare in our simulations. Our finding that bet-hedging occurs between where conditions favor infected over uninfected hosts hints at the possibility that bet-hedging could facilitate the evolution of symbiosis where benign environments transition to harsh environments. However, as our simulations also reveal, symbiosis is more often favored without the need for bet-hedging even with costs. Our results thus weaken the case that costly symbiosis in some contexts are necessarily examples of bet-hedging since symbiosis was more often favored outright than by bet-hedging.

Symbiotic interactions may play a larger role in adaptation to variable environments than previously understood, even without bet-hedging. Symbioses are known to result in novel phenotypes that allow partners to survive in harsh conditions (Moran, 2007; Oliver *et al*., 2010). Rarely do studies incorporate environmental variation and long-term fitness. We investigated the long-term effects of context-dependence in the symbiosis between *D. discoideum* and *Paraburkholderia* and found that hosts frequently benefited from symbiosis in the harshest conditions. An understanding of the ecological contexts along with long-term measures of fitness will be important for understanding the evolutionary consequences of context-dependent symbioses.

## Acknowledgements

This material is based upon work funded by the National Science Foundation under grants DEB-1753743 and IOS-1656756. We thank Tyler Larsen for figure templates. We thank Tyler Larsen, James Medina, two anonymous reviewers, and Andy Gardner for comments on the manuscript. We would also like to thank Thomas Haaland and the Strassmann/Queller lab group, especially Debbie Brock for feedback during the development and execution of this project.

## Authorship statement

TS, DQ, and JS designed the study and wrote the manuscript. TS performed the experiments and simulations and analyzed the data.

## Data Statement

Data will be archived at Dryad. Data and code for simulations and analysis will be available at www.gitlab.com/treyjscott/farmerBH.

## Notes

https://gitlab.com/treyjscott/farmerbh

### Competing Interest Statement

The authors have declared no competing interest.

## References

Bonner, J.T. (1982) Evolutionary Strategies and Developmental Constraints in the Cellular Slime Molds. The American Naturalist, 119, 530–552.

Brock, D.A., Douglas, T.E., Queller, D.C. & Strassmann, J.E. (2011) Primitive agriculture in a social amoeba. Nature, 469, 393–396.

Brock, D.A., Haselkorn, T.S., Garcia, J.R., Bashir, U., Douglas, T.E., Galloway, J., et al. (2018) Diversity of Free-Living Environmental Bacteria and Their Interactions With a Bactivorous Amoeba. Frontiers in Cellular and Infection Microbiology, 8.

Brock, D.A., Jones, K., Queller, D.C. & Strassmann, J.E. (2016) Which phenotypic traits of Dictyostelium discoideum farmers are conferred by their bacterial symbionts? Symbiosis, 68, 39–48.

Brock, D.A., Noh, S., Hubert, A.N.M., Haselkorn, T.S., DiSalvo, S., Suess, M.K., et al. (2020) Endosymbiotic adaptations in three new bacterial species associated with *Dictyostelium discoideum*: *Paraburkholderia agricolaris* sp. nov., *Paraburkholderia hayleyella* sp. nov., and *Paraburkholderia bonniea* sp. nov. PeerJ, 8, e9151.

Bronstein, J.L. (1994) Conditional outcomes in mutualistic interactions. Trends in Ecology & Evolution, 9, 214–217.

Burnham, K.P. & Anderson, D.R. (2004) Multimodel Inference: Understanding AIC and BIC in Model Selection. Sociological Methods & Research, 33, 261–304.

Buttery, N.J., Rozen, D.E., Wolf, J.B. & Thompson, C.R.L. (2009) Quantification of Social Behavior in D. discoideum Reveals Complex Fixed and Facultative Strategies. Current Biology, 19, 1373–1377.

Chamberlain, S.A., Bronstein, J.L. & Rudgers, J.A. (2014) How context dependent are species interactions? Ecology Letters, 17, 881–890.

DiSalvo, S., Haselkorn, T.S., Bashir, U., Jimenez, D., Brock, D.A., Queller, D.C., et al. (2015) *Burkholderia* bacteria infectiously induce the proto-farming symbiosis of *Dictyostelium* amoebae and food bacteria. Proceedings of the National Academy of Sciences, 112, E5029–E5037.

Douglas, A.E. (2010) The symbiotic habit. Princeton University Press.

Fey, P., Dodson, R.J., Basu, S. & Chisholm, R.L. (2013) One stop shop for everything Dictyostelium: dictyBase and the Dicty Stock Center in 2012. In Dictyostelium Discoideum Protocols. Springer, pp. 59–92.

Garcia, J.R. & Gerardo, N.M. (2014) The symbiont side of symbiosis: do microbes really benefit? Frontiers in Microbiology, 5.

Garcia, J.R., Larsen, T.J., Queller, D.C. & Strassmann, J.E. (2019) Fitness costs and benefits vary for two facultative *Burkholderia* symbionts of the social amoeba, Dictyostelium discoideum. Ecology and Evolution, ece3.5529.

Gruenheit, N., Parkinson, K., Stewart, B., Howie, J.A., Wolf, J.B. & Thompson, C.R.L. (2017) A polychromatic ‘greenbeard’ locus determines patterns of cooperation in a social amoeba. Nature Communications, 8, 14171.

Hall, D.W., Fox, S., Kuzdzal-Fick, J.J., Strassmann, J.E. & Queller, D.C. (2013) The Rate and Effects of Spontaneous Mutation on Fitness Traits in the Social Amoeba, Dictyostelium discoideum. G3 Genes/Genomes/Genetics, 3, 1115–1127.

Haselkorn, T.S., DiSalvo, S., Miller, J.W., Bashir, U., Brock, D.A., Queller, D.C., et al. (2019) The specificity of *Burkholderia* symbionts in the social amoeba farming symbiosis: Prevalence, species, genetic and phenotypic diversity. Molecular Ecology, 28, 847–862.

Haselkorn, T.S., Jimenez, D., Bashir, U., Sallinger, E., Queller, D.C., Strassmann, J.E., et al. (2021) Novel Chlamydiae and *Amoebophilus* endosymbionts are prevalent in wild isolates of the model social amoeba *Dictyostelium discoideum*. Environmental Microbiology Reports, 1758–2229.12985.

Iwai, S. (2019) Photosynthetic Endosymbionts Benefit from Host’s Phagotrophy, Including Predation on Potential Competitors. Current Biology, 29, 3114–3119.

Keeling, P.J. & McCutcheon, J.P. (2017) Endosymbiosis: The feeling is not mutual. Journal of Theoretical Biology, 434, 75–79.

Kessin, R.H. (2001) Dictyostelium: evolution, cell biology, and the development of multicellularity. Cambridge University Press.

Khojandi, N., Haselkorn, T.S., Eschbach, M.N., Naser, R.A. & DiSalvo, S. (2019) Intracellular Burkholderia Symbionts induce extracellular secondary infections; driving diverse host outcomes that vary by genotype and environment. The ISME Journal.

Kraemer, S.A. & Boynton, P.J. (2017) Evidence for microbial local adaptation in nature. Molecular Ecology, 26, 1860–1876.

Kuserk, F.T. (1980) The Relationship Between Cellular Slime Molds and Bacteria in Forest Soil. Ecology, 61, 1474–1485.

Lekberg, Y. & Koide, R.T. (2014) Integrating physiological, community, and evolutionary perspectives on the arbuscular mycorrhizal symbiosis. Botany, 92, 241–251.

Lenth, R., Singmann, H., Love, J., Buerkner, P. & Herve, M. (2018) Emmeans: Estimated marginal means, aka least-squares means. R package version, 1, 3.

Lowe, C.D., Minter, E.J., Cameron, D.D. & Brockhurst, M.A. (2016) Shining a Light on Exploitative Host Control in a Photosynthetic Endosymbiosis. Current Biology, 26, 207–211.

Meyers, A., Furtmann, C. & Jose, J. (2018) Direct optical density determination of bacterial cultures in microplates for high-throughput screening applications. Enzyme and Microbial Technology, 118, 1–5.

Miller, J.W., Bocke, C.R., Tresslar, A.R., Schniepp, E.M. & DiSalvo, S. (2020) Paraburkholderia Symbionts Display Variable Infection Patterns That Are Not Predictive of Amoeba Host Outcomes. Genes, 11, 674.

Moran, N.A. (2007) Symbiosis as an adaptive process and source of phenotypic complexity. Proceedings of the National Academy of Sciences, 104, 8627–8633.

Oliver, K.M., Degnan, P.H., Burke, G.R. & Moran, N.A. (2010) Facultative Symbionts in Aphids and the Horizontal Transfer of Ecologically Important Traits. Annual Review of Entomology, 55, 247–266.

Philippi, T. & Seger, J. (1989) Hedging one’s evolutionary bets, revisited. Trends in ecology & evolution, 4, 41–44.

Pinheiro, J. & Bates, D. (2006) Mixed-effects models in S and S-PLUS. Springer Science & Business Media.

R Core Team. (2020) R: A language and environment for statistical computing. R Foundation for Statistical Computing, Vienna, Austria.

Raper, K.B. (1937) Growth and development of Dictyostelium discoideum with different bacterial associates. Journal of Agricultural Research, 55, 289–316.

Ruokolainen, L., Lindén, A., Kaitala, V. & Fowler, M.S. (2009) Ecological and evolutionary dynamics under coloured environmental variation. Trends in Ecology & Evolution, 24, 555–563.

Sæther, B.-E. & Engen, S. (2015) The concept of fitness in fluctuating environments. Trends in Ecology & Evolution, 30, 273–281.

Seger, J. & Brockmann, H. (1987) What is bet-hedging? In Oxford surveys in evolutionary biology. Oxford University Press.

Shu, L., Brock, D.A., Geist, K.S., Miller, J.W., Queller, D.C., Strassmann, J.E., et al. (2018a) Symbiont location, host fitness, and possible coadaptation in a symbiosis between social amoebae and bacteria. eLife, 7, 25.

Shu, L., Zhang, B., Queller, D.C. & Strassmann, J.E. (2018b) Burkholderia bacteria use chemotaxis to find social amoeba Dictyostelium discoideum hosts. The ISMEJournal, 12, 1977–1993.

Slatkin, M. (1974) Hedging one’s evolutionary bets. Nature, 250, 704–705.

smith, jeff, Queller, D.C. & Strassmann, J.E. (2014) Fruiting bodies of the social amoeba Dictyostelium discoideum increase spore transport by Drosophila. BMC Evolutionary Biology, 14, 105.

Starrfelt, J. & Kokko, H. (2012) Bet-hedging-a triple trade-off between means, variances and correlations. Biological Reviews, 87, 742–755.

Suthers, H.B. (1985) Ground-feeding migratory songbirds as cellular slime mold distribution vectors. Oecologia, 65, 526–530.

Thompson, J.N. (1994) The coevolutionary process. University of Chicago Press.

Veresoglou, S.D., Johnson, D., Mola, M., Yang, G. & Rillig, M.C. (2021) Evolutionary bet-hedging in arbuscular mycorrhiza-associating angiosperms. New Phytologist, nph.17852.

Vos, M., Wolf, A.B., Jennings, S.J. & Kowalchuk, G.A. (2013) Micro-scale determinants of bacterial diversity in soil. FEMS Microbiology Reviews, 37, 936–954.

Young, I.M. (2004) Interactions and Self-Organization in the Soil-Microbe Complex. Science, 304, 1634–1637.

