## Supplemental File 1 for "Context-dependence in the symbiosis between *Dictyostelium discoideum* and *Paraburkholderia*"

**Supplemental material for Scott et al. submitted to Evolution Letters**

**Supplemental Methods**

*Continuous food simulations*

The distribution of food bacteria in nature is likely to be continuous with some patches having more or less, in contrast to our binary patches in the main text. Natural conditions also tend to exhibit temporal autocorrelations (Ruokolainen *et al.* 2009) that could impact the abundance of food. To test whether our simulation results were robust to more realistic assumptions about continuous food and correlated environmental change, we created additional simulations. Reproduction and dispersal occurred as in simulations in the main text.

To use our empirical spore production values with continuous food, we created a linear model of spore production, *y* = *af* + *b*. *y* is spore production, *a* is the slope, *f* is the amount of food, and *b* is the intercept. For each clone on each date, we determine the intercept *b* for spore production when *f* is 0 as the spore production of that clone on that date in food-poor contexts. The slope *a* is then the difference between spore production in food-rich vs food-poor contexts. This results in models that will recapitulate our empirical spore production values in food-poor (when f = 0) and food-rich (f = 1) along with continuous spore production between these extremes (Figure S4). To incorporate costs in food-rich contexts for infected hosts, we modify our linear model to include a 1- *c* term, spores = *a*(1-*c*)*f* + *b*. This results in costs only being applied to the part of the equation that is a function of food (Figure S4 with 25% cost).

To make food continuous, we assumed that food in the environment at time *t* was a function of some continuous resource *r_t_*. To model spatial variation, we drew patch resource values from a normal distribution with mean r_t_ and standard deviation *s*. At the start of individual replicates (simulations of the three strategies on identical environments), we randomly sampled the starting amount of resource *r_1_* from a standard normal distribution with mean 0 and standard deviation 1.We modeled temporal change of the resource using fractal Brownian motion (Figure S5). The amount of autocorrelation over time is controlled by varying the Hurst exponent, H (Travis 2001). When H is low, successive values of r_t_ are anti-correlated (Figure S5A). When H is 0.5, values of r_t_ are uncorrelated (Figure S5B). When H increases beyond 0.5, future values of r_t_ resemble past values (Figure S5C). H is a continuous food alternative to our v_temp_ argument in binary food environments with low H corresponding to highly variable environments and high H corresponding to less variation. For this reason, the y axes in Figures S6-8B can be loosely interpreted as the flipped version of the y axes in Figure 4.

To map the amount of resource in a patch to the amount of food, we modeled the density of a food bacterium that is best adapted to an intermediate value of the resource. In nature, this might occur when a bacterium grows best at an intermediate value of some nutrient or chemical. For example, most bacteria grow best at intermediate pHs with diminishing growth as pH becomes more basic or acidic. We assumed there is some optimal value — without loss of generality, we chose zero as the optimum — of this resource where food is maximized (at one) according to a Gaussian function, f = exp(-0.5r^2^). As the resource value moves away from the optimum, the amount of food falls off and approaches zero.

Since our binary simulations showed that a lack of food is important, we sought to similarly measure the amount of food in these continuous environments. With fractal Brownian motion, we do not explicitly control the average amount of food, but we were able to calculate average amounts of food from our simulations. In order to determine which strategies won in different food conditions, we binned average values into 20 bins and report the mean food-poor (1- the average amount of food) value in Figures S6-8. This is analogous to our mean *p* used in binary patch simulation because it gives a measure of food scarcity in the environment. Because we did not directly control the amount of food, there are different numbers of simulations in food bins. However, removal of bins with fewer simulations or changing the number of bins did not change our results.

References:

Ruokolainen, L., Lindén, A., Kaitala, V. & Fowler, M.S. (2009). Ecological and evolutionary dynamics under coloured environmental variation. *Trends Ecol. Evol.*, 24, 555–563.

Travis, J.M.J. (2001). The color of noise and the evolution of dispersal. *Ecol. Res.*, 16, 157–163.

**Supplemental Figures**

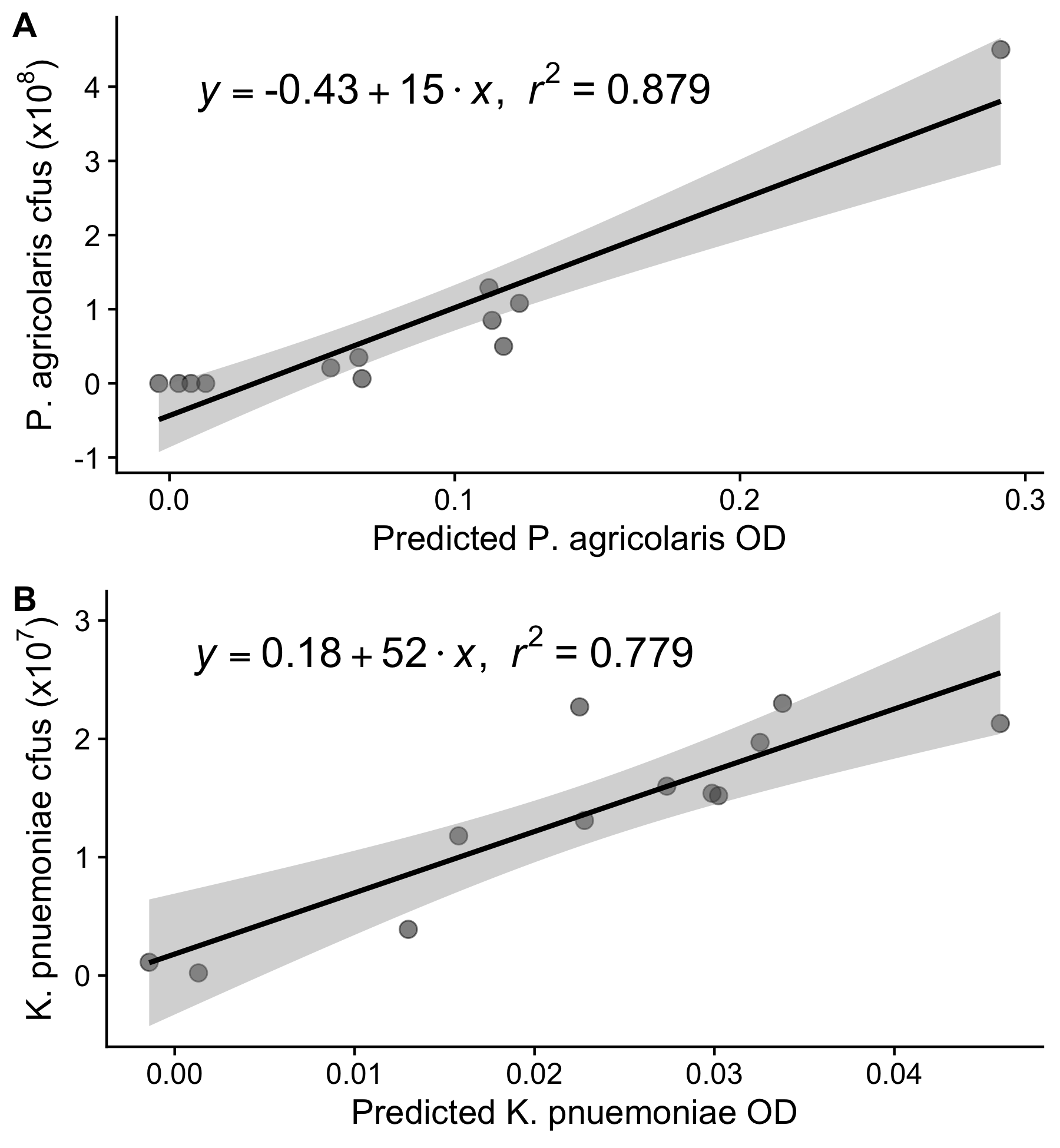

**Figure S1:** Linear regression results relating estimated optical density (predicted from standard curve) to colony counts from serial dilutions of the same *P. agricolaris* (A) and *K. pneumoniae* (B) samples.

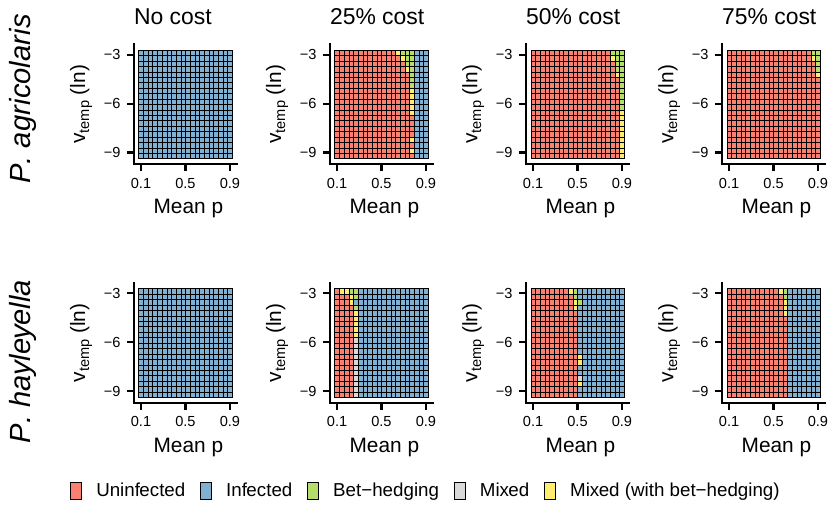

**Figure S2:** Plot of the best phenotype for a given probability of being in a no food environment and cost of having symbionts. These are the same as Figure 4 with 50% chance of successful colonization.

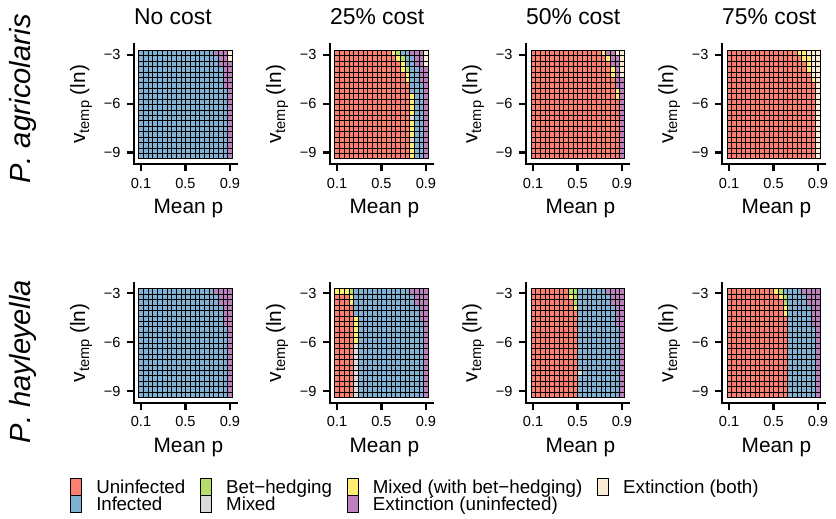

**Figure S3:** Plot of the best phenotype for a given probability of being in a no food environment and cost of having symbionts. These are the same as Figure 4 with 0.5% chance of successful colonization.

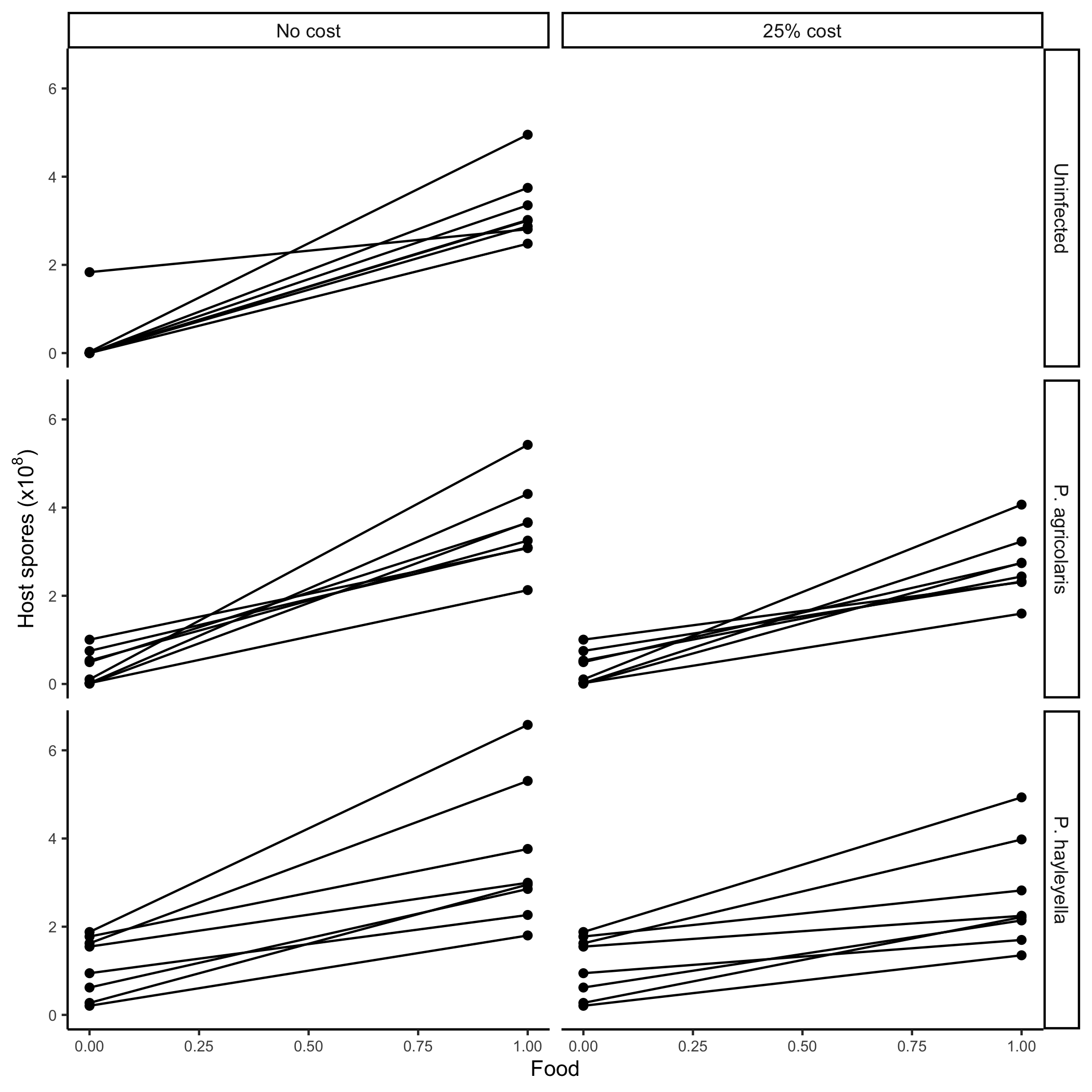

**Figure S4:** Models of continuous food without and with costs. Points show empirical spore production values from Figure 3. Lines show linear models of spore production. Costs are applied by reducing spore production when food is greater than 0.

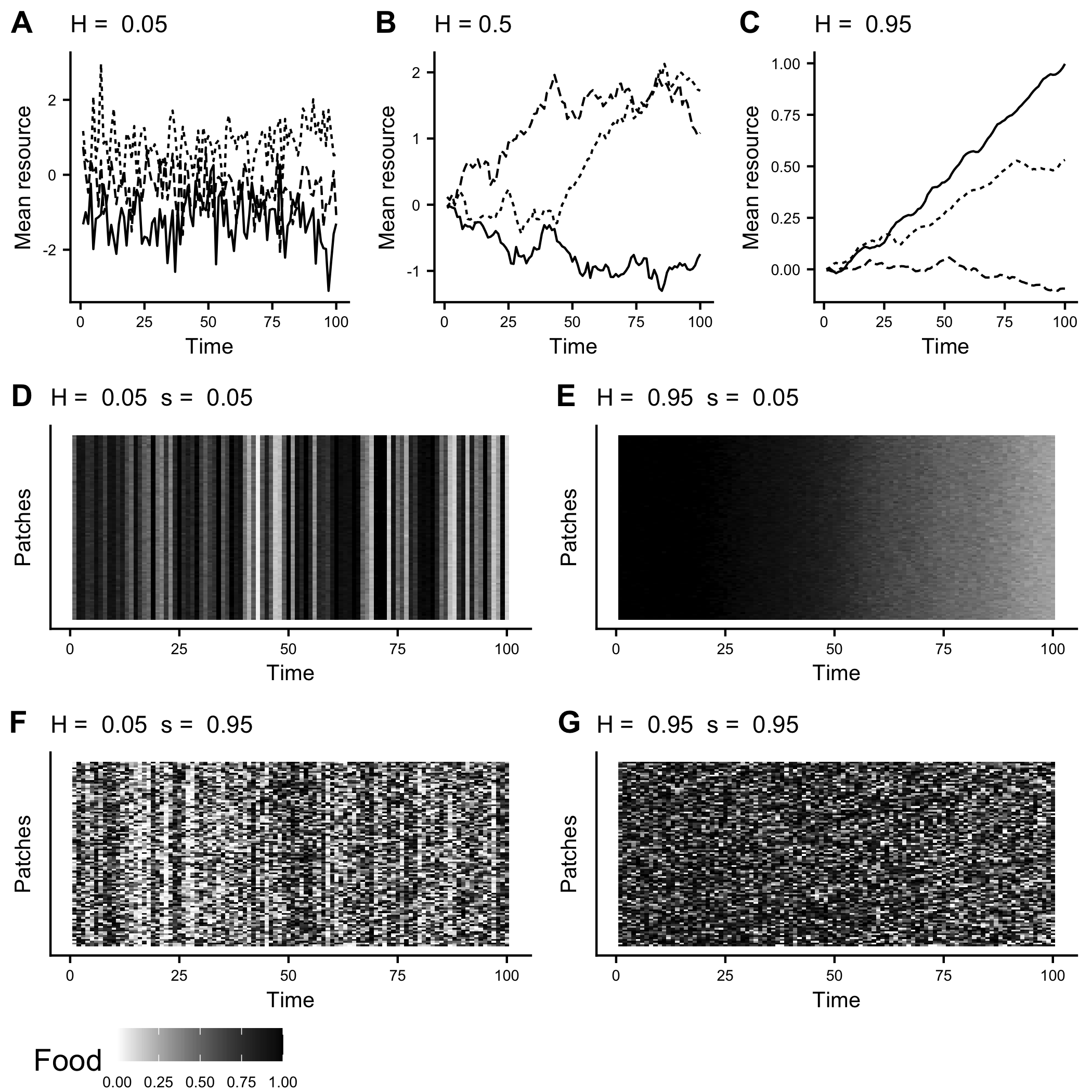

**Figure S5:** Temporally and spatially varying resource and its impact on food. (A-B) Impact of temporal variation in the mean resource for different Hurst exponents (H). (D-G) Amount of food with different H and standard deviations, *s*. D-G show how environments can vary depending on the values of H and s. In D, anti-correlated temporal variation and low spatial variation result in large differences in the amount of food across time. In E, correlated temporal variation and low spatial variation results in gradual change over time. F and G are similar to D and E, except that there is now ample variation across space.

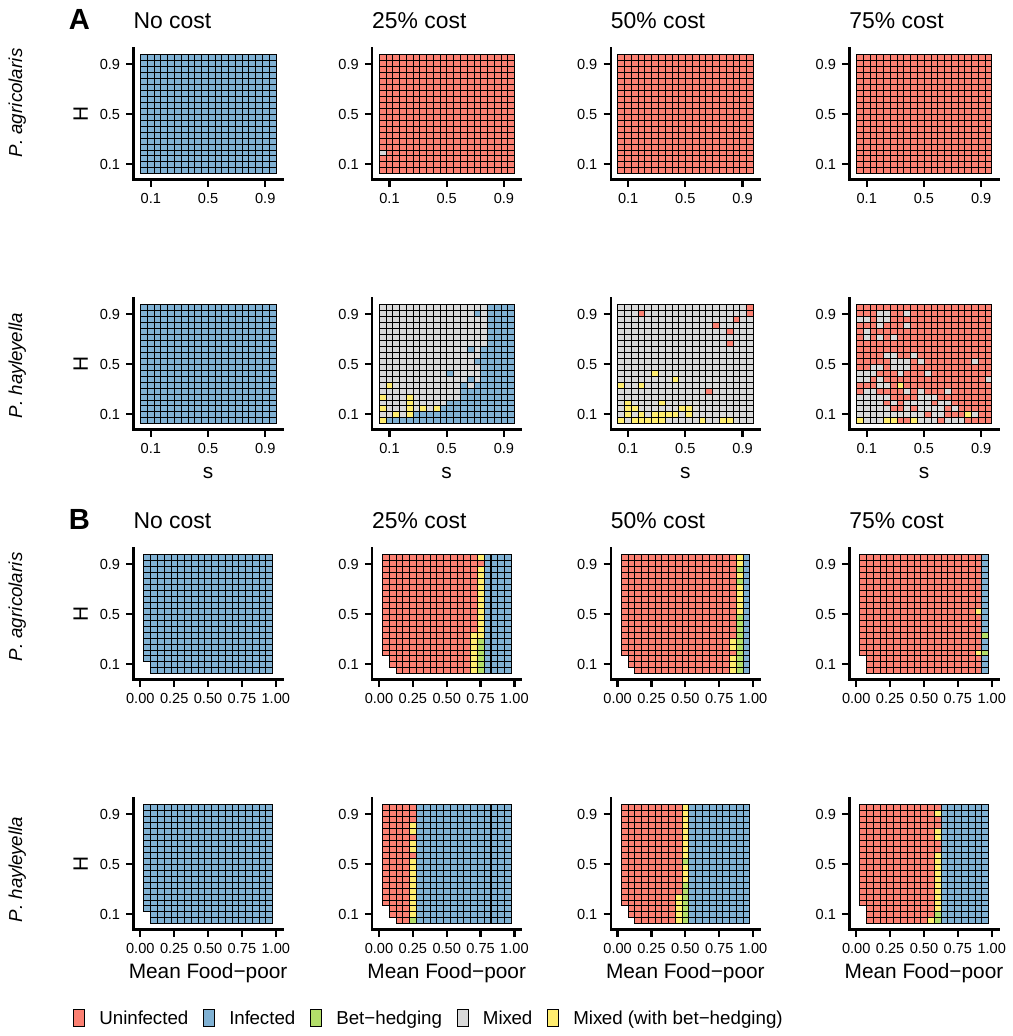

**Figure S6:** Plots of best phenotypes from continuous food simulations. (A) Best phenotypes with varying Hurst exponent (H) and standard deviation of spatial variation, *s*. (B) Best phenotypes when results were binned according to how much food was in the environment. B is analogous to Figure 4 where the y axes in B can loosely be interpreted as the flipped version of the y axes in Figure 4 in the main text. For these simulations, g = 5%.

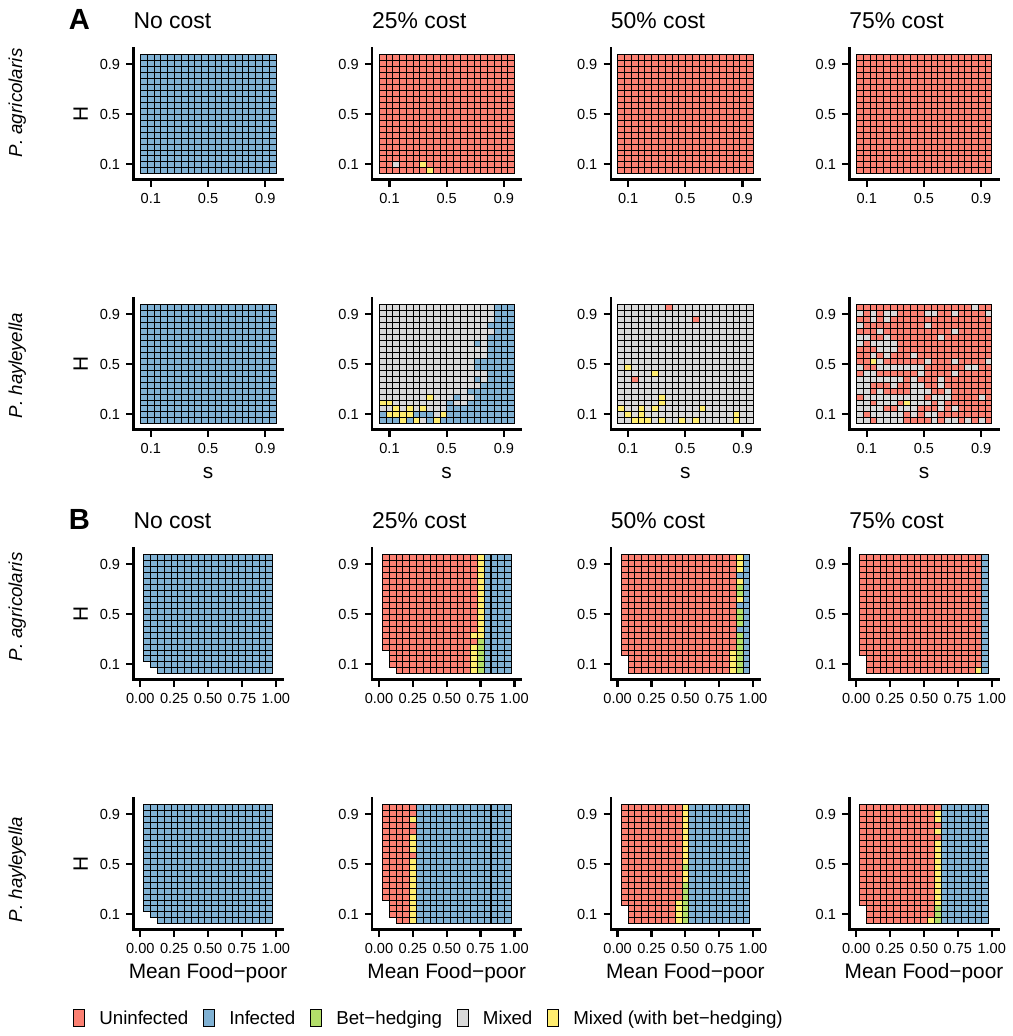

**Figure S7:** Plots of best phenotypes from continuous food simulations. (A) Best phenotypes with varying Hurst exponent (H) and standard deviation of spatial variation, *s*. (B) Best phenotypes when results were binned according to how much food was in the environment. B is analogous to Figure 4 where the y axes in B can loosely be interpreted as the flipped version of the y axes in Figure 4 in the main text. For these simulations g = 50%.

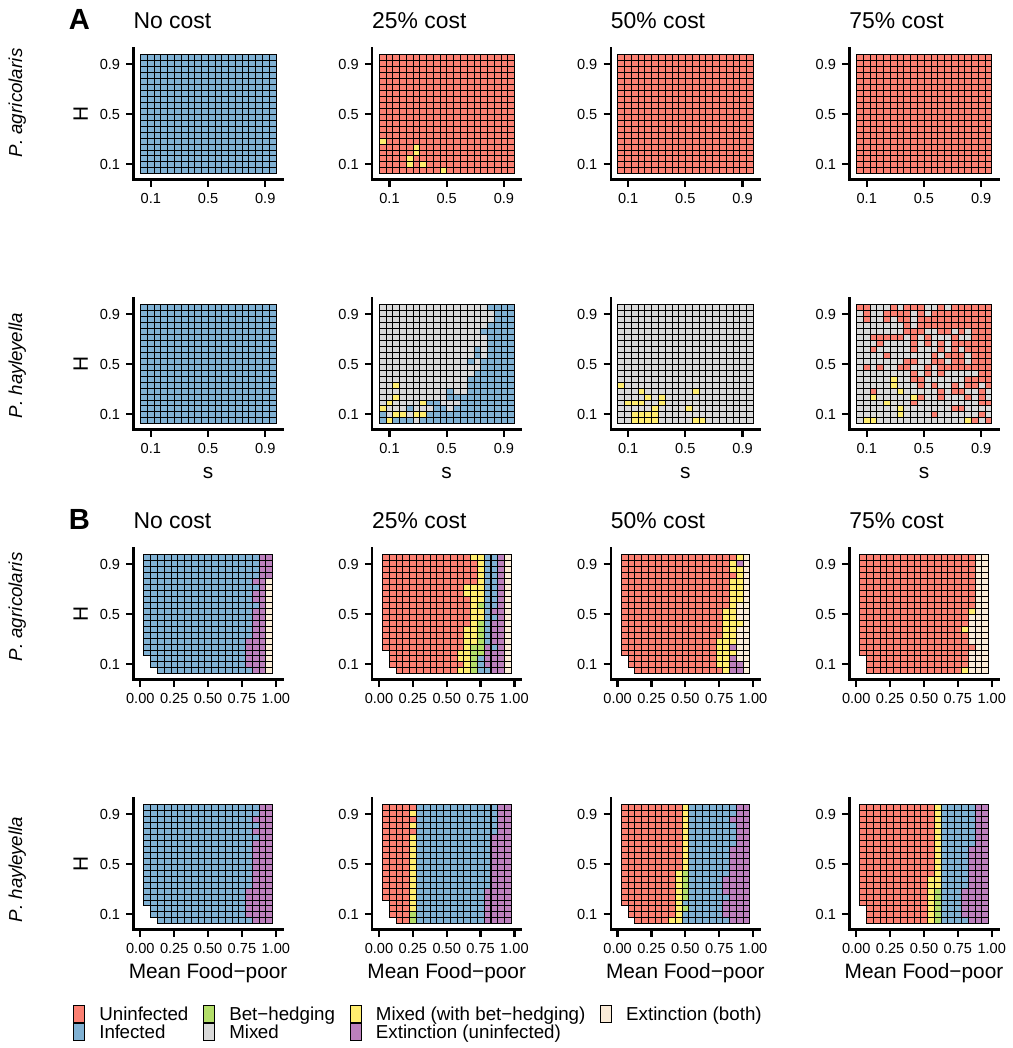

**Figure S8:** Plots of best phenotypes from continuous food simulations. (A) Best phenotypes with varying Hurst exponent (H) and standard deviation of spatial variation, *s*. (B) Best phenotypes when results were binned according to how much food was in the environment. B is analogous to Figure 4 where the y axes in B can loosely be interpreted as the flipped version of the y axes in Figure 4 in the main text. For these simulations g = 0.5%.

**Animation S1**: Example simulation with 100 rounds of growth and dispersal and a 25% cost of infection. Food and v_temp_ are the same as in the top left corner of panels in Figure 4.

**Animation S2**: Example simulation with 100 rounds of growth and dispersal and a 25% cost of infection. Food and v_temp_ are the same as the top right corner of panels in Figure 4.

**Animation S3**: Example simulation with 100 rounds of growth and dispersal and a 25% cost of infection. Food and v_temp_ are the same as the bottom left corner of panels in Figure 4.

**Animation S4**: Example simulation with 100 rounds of growth and dispersal and a 25% cost of infection. Food and v_temp_ are the same as the bottom right corner of panels in Figure 4.

**Animation S5**: Example simulation with 100 rounds of growth and dispersal and a 25% cost of infection when food is continuous (H = s = 0.05).
